# Rethinking Dormancy: Antibiotic Persisters are Metabolically Active, Non-Growing Cells

**DOI:** 10.1101/2023.12.09.570944

**Authors:** K M Taufiqur Rahman, Ruqayyah Amaratunga, Xuan Yi Butzin, Abhyudai Singh, Tahmina Hossain, Nicholas C. Butzin

## Abstract

Bacterial persisters are a subpopulation of multidrug-tolerant cells capable of surviving and resuming activity after exposure to bactericidal antibiotic concentrations, contributing to relapsing infections and the development of antibiotic resistance. We challenge the conventional view that persisters are metabolically dormant by providing compelling evidence that an isogenic population of *Escherichia coli* remains metabolically active in persistence. Our transcriptomic analysis, conducted at various time points following exposure to bactericidal concentrations of ampicillin (Amp), revealed a number of genes with differential expression over time. Some genes were consistently upregulated in Amp treated persisters compared to the untreated controls, a change that can only occur in metabolically active cells capable of increasing RNA levels. Some of these genes have been previously linked to persister cells, while others have not been associated with them before. If persister cells were metabolically dormant, we would expect minimal changes in the gene network across different time points of Amp treatment. However, network analysis revealed major shifts in gene network activity at various time points of antibiotic exposure. These findings reveal that persisters are metabolically active, non-dividing cells, thereby challenging the notion that they are dormant.

**Significance statement:** Bacterial persisters are a subpopulation renowned for their multidrug tolerance and remarkable ability to survive bactericidal antibiotic treatments; understanding their formation and long-term survival presents significant challenges. These persisters play a critical role in driving antibiotic resistance, underscoring the urgency of deepening our knowledge about them as the threat of resistance continues to escalate. Our study challenges the long-held assumption that persisters are metabolically inactive and that persisters are not as dormant as previously thought.

## Introduction

Bacteria thrive in a wide range of harsh environments, requiring them to adapt to stress by modifying their metabolic states—a phenomenon widely known as persistence [3, 4]. All free-living bacteria employ this bet-hedging strategy to survive adverse conditions [4]. Persisters are a small fraction of the bacterial population, emerging from inherent phenotypic heterogeneity rather than genetic mutations. These cells exhibit reduced cellular activity and do not divide under stress [5]. When the threat subsides, persister cells can resume growth, often leading to recurrent infections that require prolonged and repeated antibiotic therapy. For instance, *Mycobacterium tuberculosis* (which causes tuberculosis) and *Mycobacterium avium* (which causes non-contagious lung disease), treatment durations can extend to about one year for *M. tuberculosis* and up to two years for *M. avium*. Despite these prolonged treatments, relapse rates remain high [6, 7]. Recent studies indicate that persisters also possess a high mutation rate [8], which has been linked to the evolution of antibiotic resistance—a pressing public health concern. Antibiotic-resistant bacteria are now the leading cause of death worldwide, surpassing other pathogens, including those responsible for diseases like HIV or malaria [9]. Projections suggest that by 2050, antibiotic-resistant infections could result in 10 million deaths annually 2050 [10, 11] unless significant measures are taken to combat this troubling trend.

Another survival strategy bacteria employ in harsh environments is the formation of biofilms—complex communities of bacteria encased in a self-produced protective matrix that adheres to surfaces. Biofilms are notoriously challenging to eradicate due to their robust defense mechanisms and intricate architecture [12-14], often containing ∼1% persister cells [15, 16]. These persisters can endure stress (i.e., bactericidal antibiotic concentrations) enabling the biofilm to regenerate after susceptible cells are eliminated and conditions improve [17, 18]. To fully grasp how bacteria survive in natural settings, we must first understand their survival strategies in controlled laboratory conditions.

Despite being discovered over eight decades ago, the molecular mechanisms underlying it remain unclear and subject to debate. A promising method to identify the key genes and regulatory networks involved in persistence is through transcriptome profiling of persister cells using RNA sequencing (RNA-Seq), which captures the full set of mRNAs synthesized by cells at a specific moment. However, due to the low abundance of persisters and the transient nature of persister cells, studying their transcriptomic profile is highly challenging. As a result, many previous transcriptomic studies have employed sub-bactericidal concentrations or used bactericidal antibiotic concentrations for a short time [19, 20]. These approaches often include populations of short-term tolerant cells—dividing, slow-growing cells—that may not represent true persisters. It is important to note that short-term tolerance can mask persistence, and evidence shows that the two phenomena are distinct [21]. Our recent findings further demonstrate that high persistence levels do not correlate with high short-term tolerance levels [22]. Therefore, it is crucial to use bactericidal antibiotic concentrations when investigating the transcriptional regulatory mechanisms of persisters [22].

## Results and Discussion

The purpose of this work is to determine whether antibiotic persisters are truly metabolically inactive (completely dormant), as often cited, or if they actively respond to environmental stimuli by altering gene expression. An overview of the experimental plan is shown in Fig. 1. Cells were grown to stationary phase, and RNA was extracted from a persister subpopulation treated with bactericidal doses (0.1 mg/mL) of ampicillin (Amp) following dilution in fresh media (Fig. 1). The treatment duration was sufficient to eliminate short-term tolerant cells. RNA sequencing was then performed to analyze the transcriptome profile of the persister cells. The *E. coli* Dh5αZ1 strain exhibited a biphasic cell death curve, with short-term tolerant cells dying rapidly in the first phase, while the remaining persisters showed a slower, steady decline starting at ∼2 h. Rapid filtration [23, 24] and flash freezing was employed to remove dead cells and minimize RNA degradation.

**Fig. 1.**
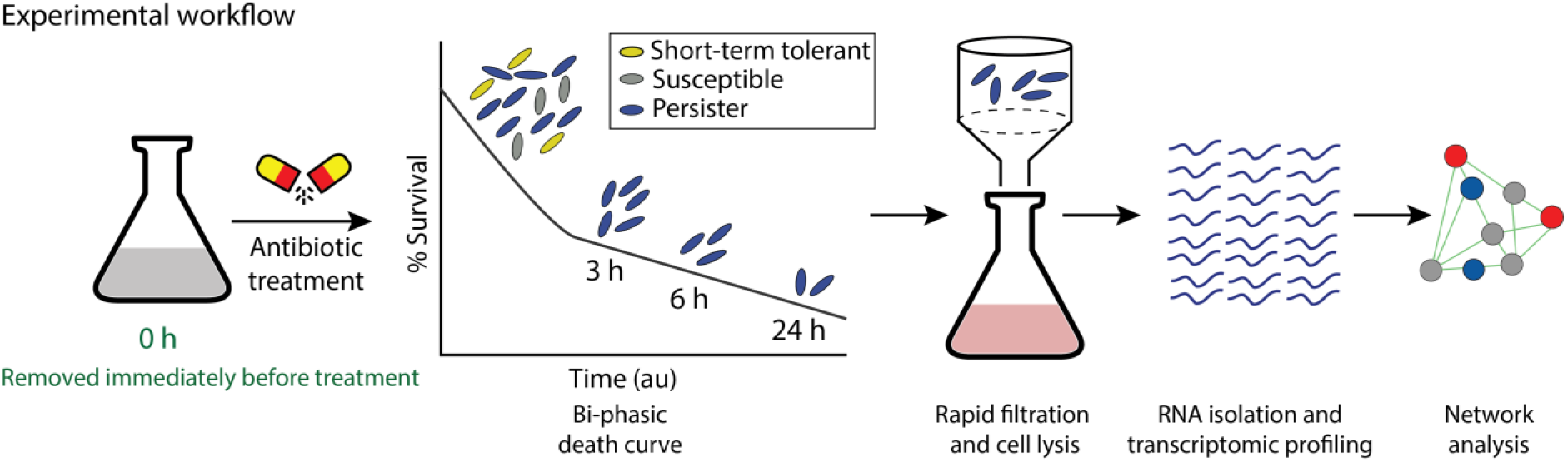
The schematic diagram of experimental design. Stationary phase cells were diluted 1:100 in pre-warmed MMB+ media and treated with 0.1 mg/mL Amp for 24 h. Samples for RNA sequencing were collected at four time points: before antibiotic treatment (0 h) and after antibiotic treatment at 3 h, 6 h, and 24 h. To minimize contamination from dead cell RNA, cells were isolated using rapid filtration. RNA extraction and sequencing were followed by gene network analysis using bioinformatics tools. Overexpression and mutational studies were then performed to validate the transcriptomic analysis.

### Contrary to the commonly held assumption of a global metabolic shutdown in persister cells, we observed differential gene expression within these cells

We compared the transcriptome profiles at different time points to investigate dynamic changes in response to bactericidal Amp treatment (0.1 mg/mL). By comparing the untreated population at 0 h with persister subpopulations at 3 h, 6 h, and 24 h post-treatment, we identified shifts in gene expression. Specifically, 378 genes were differentially expressed at 3 h, 1478 at 6 h, and 310 at 24 h (Fig. 2a). Interestingly, these genes exhibited diverse patterns of upregulation and downregulation at 3 h, 6 h, and 24 h of treatment (Fig. 2a). Contrary to the commonly held hypothesis of a global metabolic shutdown in persister cells, which would predict no gene upregulation, we observed significant upregulation of several genes. The heatmap clearly showed distinct expression patterns across all gene sets and time points (Fig. 2b). This led us to focus on a specific set of genes with similar expression profiles. Notably, 27 genes were consistently upregulated across all time points (Fig. 2c). Of these, 8 were hypothetical genes (Table S1). The upregulation of hypothetical genes, which have not been previously linked to antibiotic survival, was particularly intriguing. We hypothesize that these genes may play a critical role in the adaptive response of persister cells, potentially contributing to their survival under antibiotic stress.

**Fig. 2.**
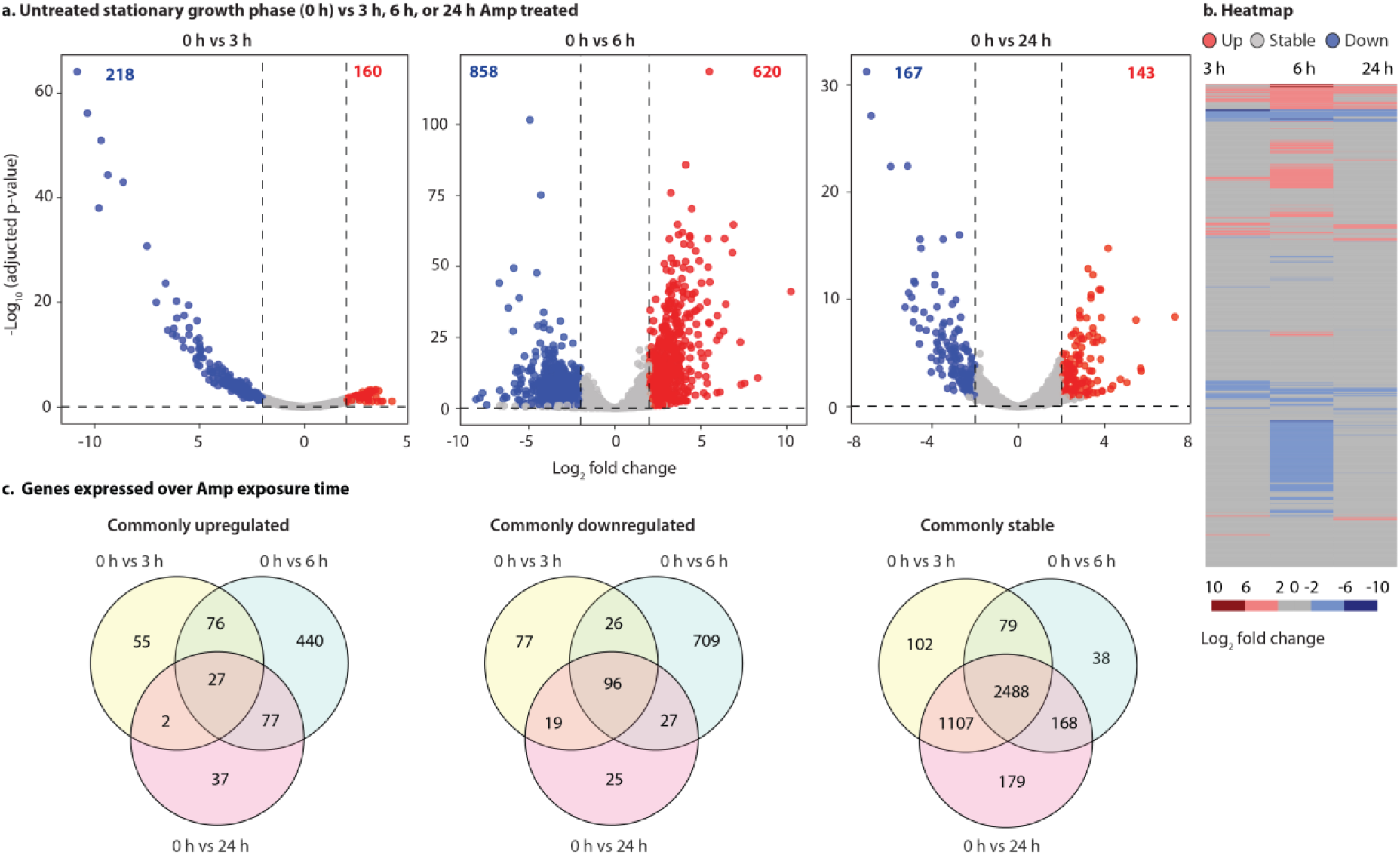
Comparative transcriptome analysis of the population before and after antibiotic treatment reveals that gene expression changes substantially over time. **a-c**: The 0 h sample is stationary-phase cultures before treatment. **a**. Volcano plots comparing gene expression at 0 h (non-persister population) with persister populations at later time points. **b**. Heatmaps of gene expression shows distinct changes in expression levels throughout treatment compared to untreated 0 h. Each row represents a gene, and each column corresponds to a treatment time point. The color intensity indicates expression levels, with red signifying high expression and blue signifying low expression. **c**. Venn diagrams depicting transcriptome profiles, where differentially expressed genes are defined by an adjusted p-values <0.1 and a fold change greater than 4 (these parameters are consistent for all subsequent figures, except for the dataset from log-phase cells, where the p-value threshold was set at 0.05).

The majority of genes (86%, 63%, and 90% at 3, 6, and 24 hours of antibiotic treatment, respectively) showed little to no change in expression compared to untreated stationary-phase cultures (Table S2). However, a notable proportion of genes (14%, 37%, and 10% at 3, 6, and 24 hours, respectively) exhibited changes in expression. These results underscore an important aspect of persister cell activity. If cells were metabolically inactive, all gene expressions would be uniformly downregulated. However, most genes continue to be expressed at levels similar to those before antibiotic treatment. Furthermore, some genes are upregulated or downregulated, indicating that gene expression in persisters is dynamic and that these cells are actively responding to their environment. For instance, after 3 h of antibiotic treatment, 160 genes are upregulated, representing ∼3.7% of the total genes in the genome.

To determine whether the upregulation of these genes was specific to the antibiotic used, we tested ciprofloxacin (Cip), which has a different mode of action than Amp. While Amp targets cell wall formation [25], Cip disrupts DNA synthesis [26]. Of the 27 genes upregulated during the 3 h, 6 h, and 24 h Amp treatment, 22 genes were also commonly upregulated after 6 h of bactericidal Cip treatment (0.01 mg/mL). This indicates a common pattern of gene activation across different antibiotics (Fig. S1). A useful way to interpret our transcriptional analysis is by examining genes already linked to persistence, thus building on insights from previous studies on persister cells and transcriptomics [25-34]. Previous studies have identified several persister-related genes, including *oxyR, dnaK, sucB, relA, rpoS, clpB, mqsR*, and *recA* [27]. We found that these previously identified persister-related genes exhibited varying responses at different time points with Amp (Table S1) and Cip treatments (Fig. S1). While all of these genes were upregulated at some point, none maintained consistent upregulation throughout the 3 h, 6 h, and 24 h treatment periods.

### Network analysis revealed significant shifts in gene network responses at various time points during antibiotic treatment

Using a cluster-centric top-down approach, we analyzed changes in gene network response and identified key network hubs at 3 h, 6 h, and 24 h of Amp treatment. While studying protein and gene networks—along with their interconnections (network topology)—has inherent limitations, such as constraints in predicting interactions and the variability of dynamic changes across cell types [35, 36], this method remains a powerful tool for uncovering interactions that are difficult to detect through other approaches. The rise of systems biology and mathematical modeling, including graph theory and neural networks, has enabled the integration of core cellular components (genes, RNAs, proteins, etc.) into a unified, complex network [37]. Applying this systematic framework, we gained critical insights into the network responses of differentially expressed genes at various time points (Fig. 3a). We constructed gene networks and identified major gene clusters using the Molecular Complex Detection (MCODE) method [38], which pinpoints highly dense clusters within a network. The gene network response fluctuated significantly across different time points, with noticeable changes in edges (connections) (Fig. 3b i) and associated nodes (genes) forming gene clusters (Fig. 3b ii and Fig. S2 a-c). The number of upregulated and downregulated genes were different at 3 h, 6 h, and 24 h (Fig. 3b ii).

**Fig. 3.**
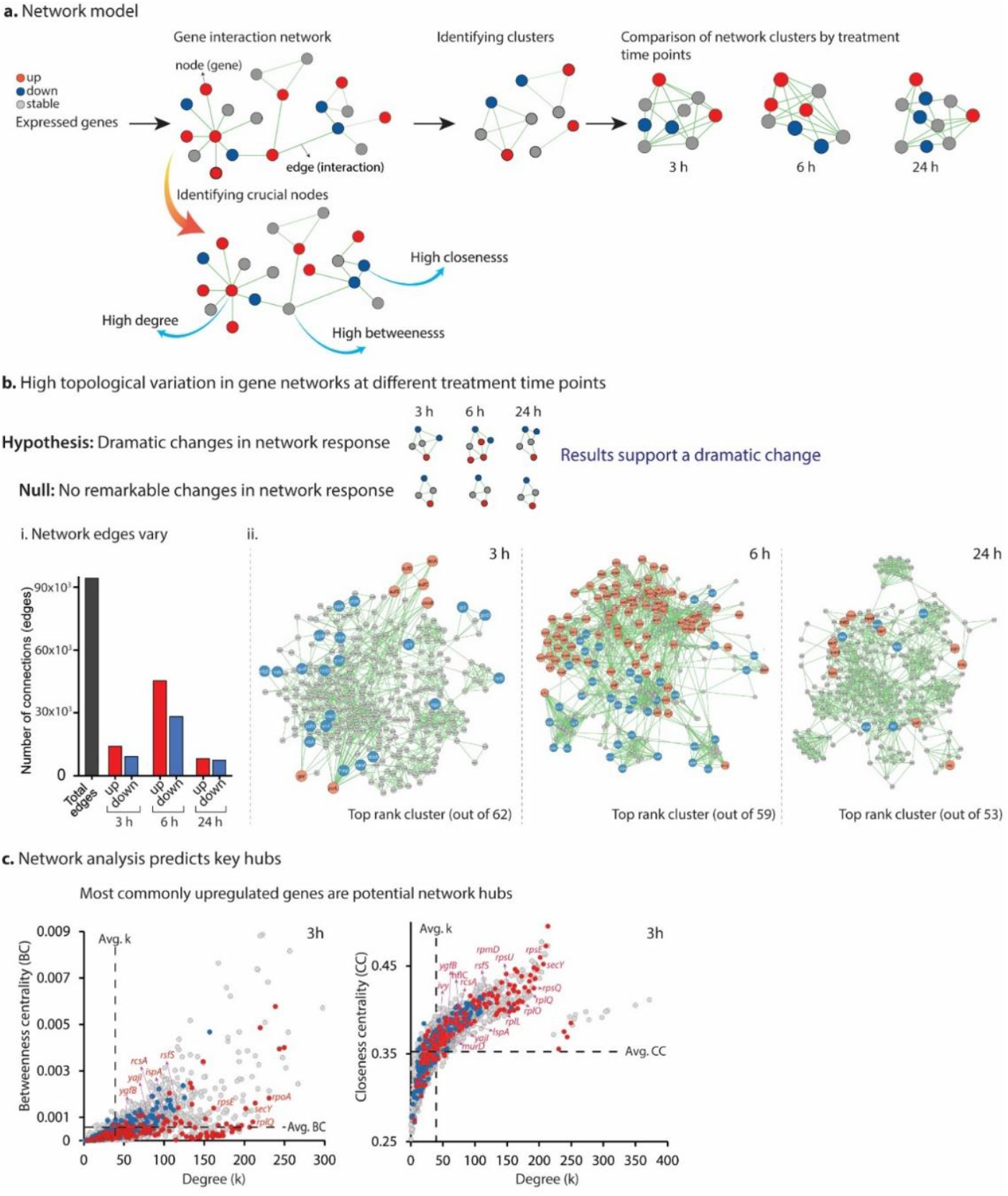
a. Interaction Network Model: In this model, each gene is represented as a node, with interactions between genes illustrated as edges. The network was analyzed using MCODE to identify gene clusters at various time points, which were subsequently compared to highlight differences in gene network responses. Key nodes within the network were pinpointed through centrality analyses encompassing node degree, betweenness, and closeness. **b. Dynamic Topological Variation:** The network analysis reveals significant topological changes in gene networks across different treatment time points. **i. Fluctuations in Edges:** A comparison of the number of edges within the network reveals fluctuations relative to the total edge count. **ii. Top-Ranking Clusters:** The analysis identified distinct clusters at 3 h (total clusters: 62), 6 h (total clusters: 59), and 24 h (total clusters: 53). The remaining clusters for each time point are presented in Fig. S2 a-c. The total number of nodes and edges varied substantially at 3 h, 6 h, and 24 h. **c. Degree Centrality:** Degree centrality represents the distribution of node connections (degrees) within the network, offering insights into its structure and revealing important characteristics related to connectivity and overall network behavior. Betweenness centrality (often referred to as a bottleneck) quantifies how frequently a node appears on the shortest paths between pairs of other nodes in the network [1], while closeness centrality measures how quickly a node can interact with others [2]. Network analysis predicts key hubs through Degree (k) vs. Betweenness Centrality (BC), and Degree (k) vs. Closeness Centrality (CC) analyses. At 3 h, commonly upregulated genes were identified as potential network hubs, with similar findings for centrality analyses at 6 h and 24 h (Fig. S3 i-ii).

Network analysis revealed a dramatic change in gene network response (Fig. 3b and Fig. S2 a-c), indicating that the overall connectivity of a gene plays a major role in regulating its expression. This suggests that multiple genes or pathways are likely to contribute to bacterial survival under stress. This happens when survival is driven by a “one-to-many” gene interaction, rather than a simple “one-to-one” interaction. Cell-state transitions and network switching facilitate this process. Much like the dynamic network switching seen in cancer cell metastasis [39, 40]. The dynamic network switching of genes may allow reversible phenotypic plasticity (cell-state transition), which causes resistance to drugs and benefits cell survival.

Typically, under normal conditions, network clusters undergo minimal to no significant alterations. However, in response to external stress (such as Amp exposure), substantial changes in gene expression and network reconfiguration are highly probable. These observations reinforce the concept that persister cells are not in a static or inactive state but are actively responding to their environment. This conclusion aligns with transcriptomic data, network analysis, and Darwin’s Theory of Evolution, which posits that organisms better adapted to their environment have a higher likelihood of survival. Persisters that can dynamically adjust their networks in response to stress are more likely to endure over time.

### Network centrality analysis reveals potential key hubs within persister gene networks

Our objective was to identify key gene hubs that are important for surviving antibiotics. We assessed each gene’s network centrality—encompassing metrics such as degree, betweenness, and closeness—to elucidate the characteristics of genes within their respective networks. Originally developed to analyze social networks, classic network centrality [41] has been adapted for biological contexts to elucidate the hierarchy of nodes and edges within complex networks. This analysis provides insights into a gene’s significance in a central network based on its connectivity and the flow of information. A high centrality score indicates that a node possesses a greater-than-average number of connections within the network. A key aspect of our analysis was contrasting the centrality scores of differentially expressed genes with the average centrality score of all genes within the network. Notably, 23%, 21%, and 21% of expressed genes exhibited high betweenness centrality scores at 3 h, 6 h, and 24 h, respectively (Fig. 3c; Fig. S3 i-ii). Similarly, when considering closeness centrality, 33%, 22%, and 31% of genes displayed high scores at 3 h, 6 h, and 24 h, respectively (Fig. 3c; Fig. S3 i-ii). These results indicate that a substantial portion of genes maintain a central position regarding their ability to interact with other genes within the network rapidly. A robust node (genes) plays a significant role, as this node is minimally affected by disruption of other nodes in a network [42], and this might be the central gene for bacterial persisters. What is particularly intriguing is the overlap between genes showing high betweenness and closeness centrality at all time points. Most of the commonly upregulated genes (excluding the hypothetical genes) showed high betweenness and closeness centrality, suggesting their importance to persister survival. This observation suggests that these genes might play critical regulatory roles early in the process, thereby contributing to the long-term survival of persister cells. It is not surprising that 8 of the 27 commonly upregulated genes exhibited lower centrality scores than the average for all genes, given that these genes are classified as hypothetical.

### The network analysis of transcriptional data revealed gene clusters with potential biological functions

We explored the roles of various genes at different time points by integrating biological interpretations and functional gene groupings using ClueGO, a Cytoscape plug-in [43, 44] (Table S3 and Fig. S3). Our focus was on genes associated with oxidative stress, cellular processes, and antibiotic responses. We found gene (both upregulated and downregulated) clusters with significant fluctuation at different time points (Table S3), suggesting a potential shift in the cellular dynamics or adaptation mechanisms. We identified clusters of both upregulated and downregulated genes that exhibited meaningful fluctuations at different time points (Table S3), indicating potential shifts in cellular dynamics or adaptive mechanisms. Notably, 6 h had the highest number of upregulated genes linked to oxidative, cellular, and antibiotic responses, but no such genes were upregulated at 24 h. This absence suggests that either other genes became more critical at 24 h or that the oxidative, cellular, and antibiotic response proteins produced at 6 h remained intact and functional, negating the need for further expression by 24 h.

### Transcriptional analysis of untreated log phase cells compared to different hours of Amp treatment shows that cells are actively producing RNA

In the RNA-seq experiment described above, we opted to compare stationary phase cells (0 h) to persisters for technical reasons. Accurate RNA-seq comparisons between treatments depend on proper normalization, which is most effective when the transcriptomic profiles of the samples being compared are similar. Based on our prior research [34] and the understanding that stationary phase cultures are enriched with persisters [45-48], we reasoned that stationary phase cells would serve as a more appropriate control than log phase cultures. Despite the challenges associated with using log phase cells (0 h) as a control, we conducted a second RNA-seq experiment comparing Amp-treated persister cells to log phase cells (0 h). If persister cells are metabolically active, then some genes should be upregulated while in the persister state.

For the second RNA-seq experiment, we adhered to the same procedure for collecting cell samples and harvesting RNA, but cells were grown to mid-log phase (∼OD 0.5) instead of stationary phase before antibiotic treatment. As anticipated, the transcriptome of persister cells differed markedly from that of log-growing cells (0 h log). This substantial divergence rendered the median of ratios normalization method used in DESeq2 unsuitable for this analysis. The median of ratios method assumes that most genes remain unchanged when comparing samples [49], an assumption that does not apply here due to the extensive global shift in gene expression. An alternative approach for samples exhibiting such a global shift is to normalize using a set of “housekeeping” genes with DESeq2. Housekeeping genes are typically expressed at relatively stable levels across various conditions; however, we could not utilize this approach because there are no well-established housekeeping genes for our specific conditions. Genes commonly used as housekeeping genes, such as *dnaK* and *recA*, could not be used because they have previously been associated with persister survival [27].

Unable to select the most suitable normalization method for our data, we applied four different approaches (Fig. 4a). Given the insufficient evidence to favor one normalization method over another, drawing quantitative conclusions from this analysis could lead to misinterpretation. Nonetheless, a key qualitative result stands out: across all normalization methods, many genes are upregulated in persister cells compared to log phase cells (Fig. 4b).

**Fig. 4.**
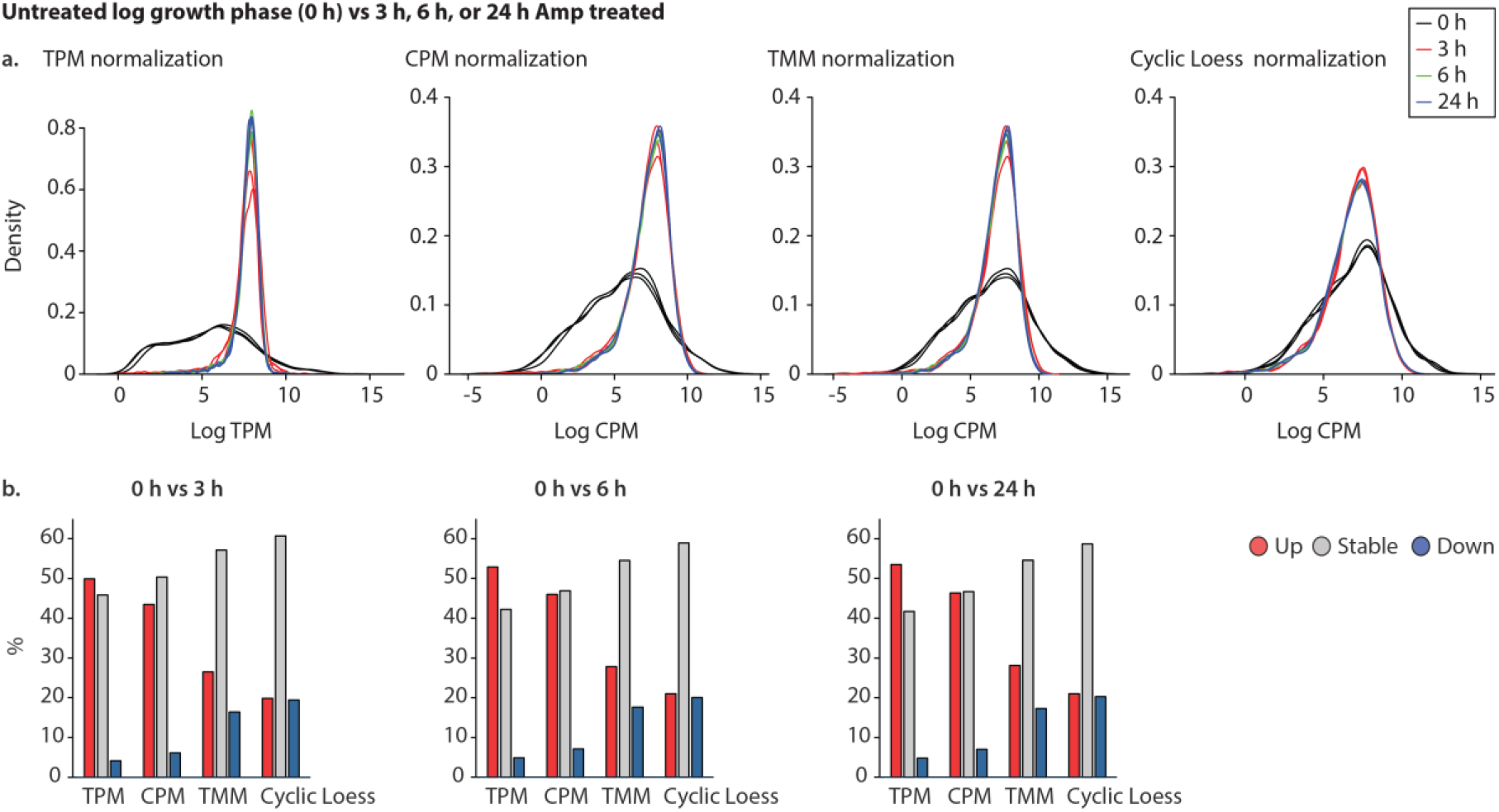
Untreated 0 h comes from mid-log phase growing cultures (OD ∼0.5). *i*. Four methods of normalization were applied, consistently showing that antibiotic-treated samples differed from the 0 h control. Transcript Per Million (TPM), Counts Per Million (CPM), Trimmed Mean of M-values (TMM), and Cyclic Loess normalization. Both TPM and CPM normalize by sequence depth, with TPM also taking gene length into account. In contrast, TMM and Cyclic Loess address compositional differences between samples, making them better equipped to handle outliers and global shifts in gene expression. Furthermore, Cyclic Loess specifically adjusts for the distribution of expression values across different samples. ***ii***. Regardless of the normalization method, several genes were consistently upregulated.

### Persister cells are metabolically active, non-growing cells that cannot be accurately described as dormant

It is common to define persisters as dormant and metabolically inactive. A leader in the persister field has stated, “These persister cells arise due to a state of dormancy, defined here as a state in which cells are metabolically inactive,” (Review article) [15]. While this assumption has underpinned much of the research on persisters, we challenge the use of the term “dormancy” when it implies that these cells are metabolically inactive or nearly inactive—failing to express new genes, produce new proteins, or respond to their environment. Although persisters do not grow [50], our findings indicate that non-growing cells can remain metabolically active and thus cannot be considered dormant. Drawing on previous work and recent studies) [34], we argue that persisters exhibit altered cellular activity rather than exist in a dormant state. This is akin to the difference between cells in stationary phase and those in the exponential phase, as both are active but possess distinct metabolic profiles. We have previously demonstrated that specific genes are upregulated in persisters compared to pre-treated (0 h) [34]. Conversely, some researchers strongly contend that the expression changes of a limited number of genes indicate that these cells are largely dormant. This perspective led to the investigation described in this manuscript, where we examined whether cells are completely, mostly, or not at all dormant. In this section, we present four lines of evidence demonstrating that persister cells are metabolically active:

#### 1. Gene Expression Patterns

If persister cells were truly dormant, we would expect all genes to be downregulated during their persistence compared to non-persisters. Our findings indicate that during the persister state, a significant proportion of genes are not downregulated; in fact, many maintain their expression levels relative to stationary phase cells.

#### 2. Upregulation of Genes

If persisters were completely inactive, we would not expect any genes to be upregulated. However, after 3 h of antibiotic treatment, we identified 160 upregulated genes compared to untreated stationary phase cells, accounting for approximately 3.7% of the genome.

Only metabolically active cells can produce new mRNA. In contrast, inactive cells would lack the capacity to synthesize new RNA or proteins for survival and would rely solely on pre-existing resources. Cellular maintenance for log and stationary cells requires substantial energy [51]. While we lack precise estimates of the energy expenditure of persister cells, it is reasonable to assume that, under the extreme stress of antibiotic treatment, maintenance constitutes a significant metabolic cost. For instance, *E. coli* has been shown to increase energy usage in response to heat [52].

While metabolically inactive cells may survive briefly without energy allocated to growth, persisters can endure much longer. Throughout this long period, persister cells must actively maintain their structural integrity and membrane potential. Cells that do not maintain their membrane potential over time will die [53, 54]. This duration represents a significant period for bacteria to survive, and the assumption that there is no change in metabolic activity throughout this time appears highly implausible for intact cells. While minimal to no metabolic activity characterizes bacterial spores, this does not apply to persister cells. Spores are so metabolically inactive that their reactivation upon germination is often described as a return to life [55-58]. Although persisters are resilient cells, they are not spores.

#### 3. Gene Interaction Network Analysis

If the cells were dormant, the interaction model should display similar profiles at 3 h, 6 h, and 24 h compared to stationary phase untreated cells. Our analysis revealed significant differences showing that metabolism of persister cells is not static.

#### 4. Supporting Evidence from Other Studies

Pu and colleagues have showed that during β-lactam antibiotic treatment, persister cells upregulate the expression of the multidrug efflux gene *tolC* [20]. This adaptation enables persisters to expel antibiotics from their cells, thereby enhancing their survival [20]. Only metabolically active cells can produce new *tolC* mRNA and new proteins.

Each line of evidence presented in this study has its own limitations, and when considered in isolation, it might seem reasonable to cling to the traditional view of persister cells. However, when these findings are examined collectively, findings support that persister cells are metabolically active. This study challenges the conventional perspective of persister cells as metabolically dormant, instead revealing that they are metabolically active, non-growing cells.

## Methods

### Microbial strains and media are described in Table S5

#### Persister assay

The assay was done as previously described [22, 34]. Cells were treated using 10X the MIC (0.01 mg/ml) for Amp and 100X the MIC (0.00001) for Cip at 37 °C and shaken at 250 rpm as previously described for this strain [59]. For the time-kill curve, antibiotic treatment was extended to 24 h, and the number of persisters was recorded at 3 h, 6 h, and 24 h.

#### RNA sample preparation, sequencing, and analysis are described in Table S6

We used our previously published data (NCBI GEO: GSE156896) [34] and new data: PRJNA1067386 and GSE278938.

### Network construction and data analysis are described in Fig. S2

## Ethical Approval

Not required

## Authors’ contributions

K.R., T.H., and R.A., performed the experiments and ran data analysis. A.S. helped analyze data and direct the project. K.R. analyzed the RNA-seq data from stationary growth phase, and X.Y.B analyzed the RNA-seq data from log growth phase. N.C.B. and T.H. formalized the research concept, designed the experiments, and planned and directed the project. All authors contributed to discussing and editing the manuscript.

## Acknowledgments

We extend our gratitude to Dr. Xijin Ge for his insightful advice and discussions on differential expression analysis. Additionally, we gratefully acknowledge the Bioconductor community forum for their helpful insights on RNA-seq normalization.

## Data availability

All the data that supports the findings of this study are included in this published article (and its supplementary information files). Also, any additional data is available from the corresponding author upon request.

