## Supplemental Files for "Rethinking Dormancy: Antibiotic Persisters are Metabolically Active, Non-Growing Cells"

**Supplement files (Figures and Tables)**

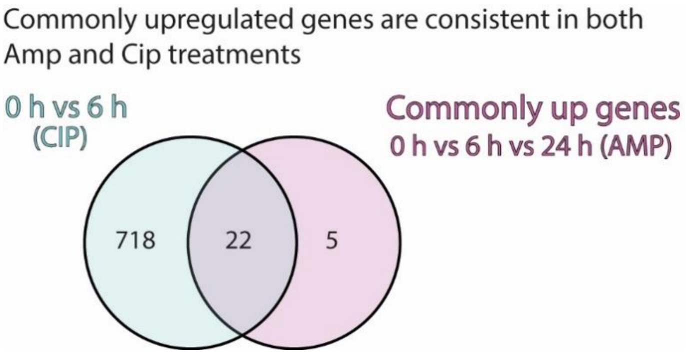

**Fig. S1.** Comparative analysis of upregulated genes in ampicillin (Amp) and ciprofloxacin (Cip) treatments (10 μg/mL) for 6 h. There were 22 genes commonly upregulated in both Amp and Cip treatments, meaning that the differential expression of these genes is not specific to Amp treatment.

**a.**

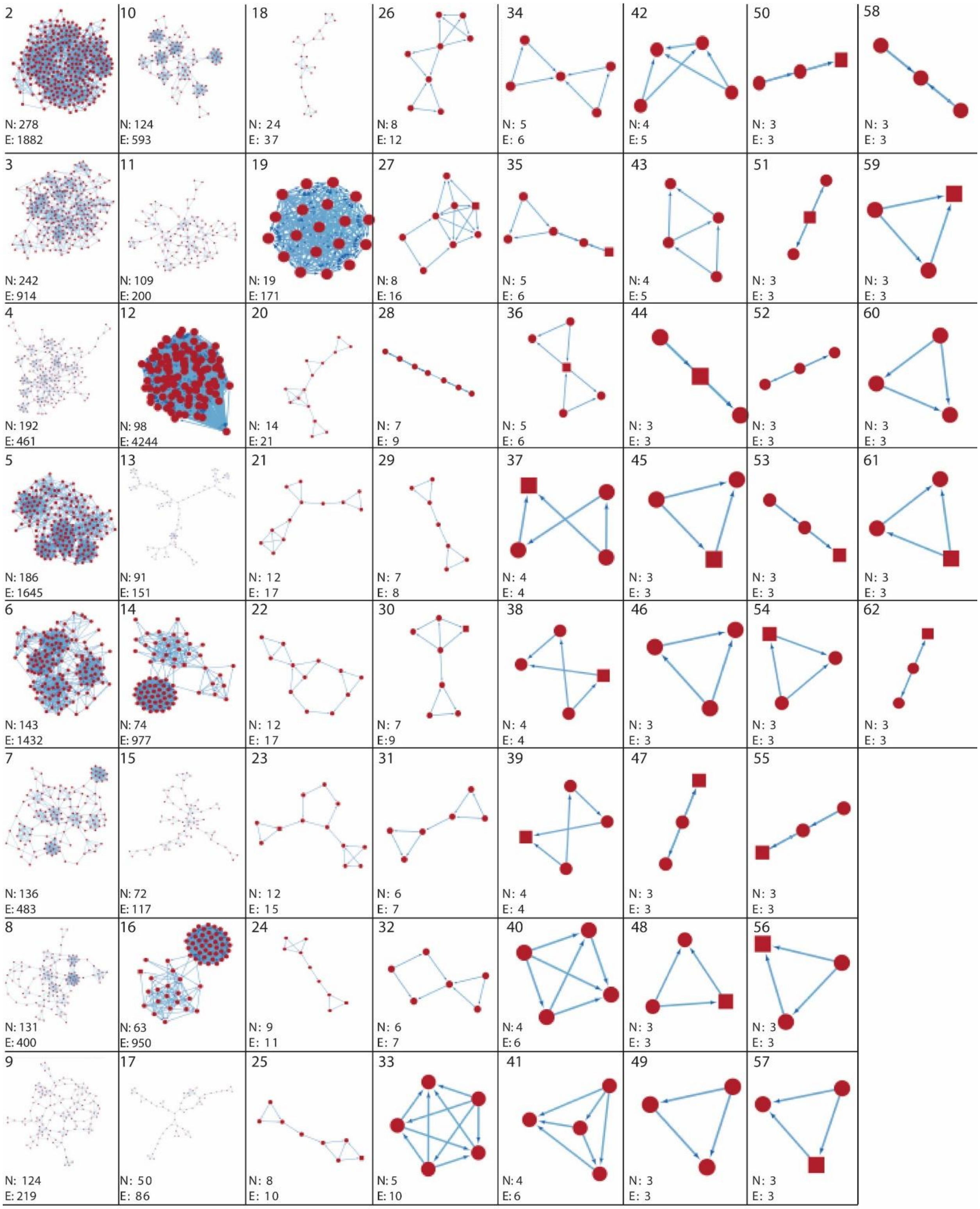

2
N: 278
E: 1882
3
N: 242
E: 914
4
N: 192
E: 461
5
N: 186
E: 1645
6
N: 143
E: 1432
7
N: 136
E: 483
8
N: 131
E: 400
9
N: 124
E: 219
10
N: 124
E: 593
11
N: 109
E: 200
12
N: 98
E: 4244
13
N: 91
E: 151
14
N: 74
E: 977
15
N: 72
E: 117
16
N: 63
E: 950
17
N: 50
E: 86
18
N: 24
E: 37
19
N: 19
E: 171
20
N: 14
E: 21
21
N: 12
E: 17
22
N: 12
E: 17
23
N: 12
E: 15
24
N: 9
E: 11
25
N: 8
E: 10
26
N: 8
E: 12
27
N: 8
E: 16
28
N: 7
E: 9
29
N: 7
E: 8
30
N: 7
E:9
31
N: 6
E: 7
32
N: 6
E: 7
33
N: 5
E: 10
34
N: 5
E: 6
35
N: 5
E: 6
36
N: 5
E: 6
37
N: 4
E: 4
38
N: 4
E: 4
39
N: 4
E: 4
40
N: 4
E: 6
41
N: 4
E: 6
42
N: 4
E: 5
43
N: 4
E: 5
44
N: 3
E: 3
45
N: 3
E: 3
46
N: 3
E: 3
47
N: 3
E: 3
48
N: 3
E: 3
49
N: 3
E: 3
50
N: 3
E: 3
51
N: 3
E: 3
52
N: 3
E: 3
53
N: 3
E: 3
54
N: 3
E: 3
55
N: 3
E: 3
56
N: 3
E: 3
57
N: 3
E: 3
58
N: 3
E: 3
59
N: 3
E: 3
60
N: 3
E: 3
61
N: 3
E: 3
62
N: 3
E: 3

**b.**

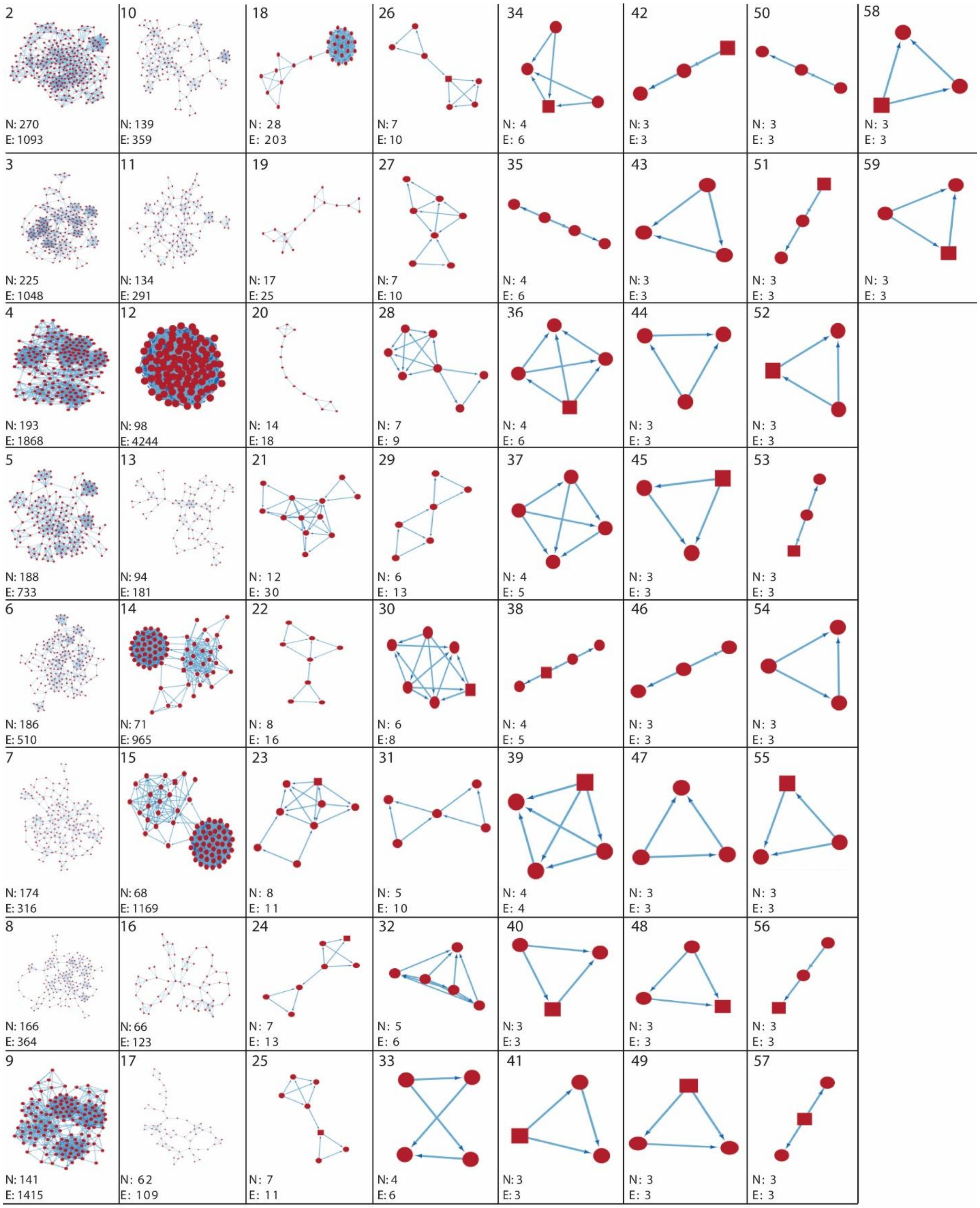

2
N: 270
E: 1093
3
N: 225
E: 1048
4
N: 193
E: 1868
5
N: 188
E: 733
6
N: 186
E: 510
7
N: 174
E: 316
8
N: 166
E: 364
9
N: 141
E: 1415
10
N: 139
E: 359
11
N: 134
E: 291
12
N: 98
E: 4244
13
N: 94
E: 181
14
N: 71
E: 965
15
N: 68
E: 1169
16
N: 66
E: 123
17
N: 62
E: 109
18
N: 28
E: 203
19
N: 17
E: 25
20
N: 14
E: 18
21
N: 12
E: 30
22
N: 8
E: 16
23
N: 8
E: 11
24
N: 7
E: 13
25
N: 7
E: 11
26
N: 7
E: 10
27
N: 7
E: 10
28
N: 7
E: 9
29
N: 6
E: 13
30
N: 6
E: 8
31
N: 5
E: 10
32
N: 5
E: 6
33
N: 4
E: 6
34
N: 4
E: 6
35
N: 4
E: 6
36
N: 4
E: 6
37
N: 4
E: 5
38
N: 4
E: 5
39
N: 4
E: 4
40
N: 3
E: 3
41
N: 3
E: 3
42
N: 3
E: 3
43
N: 3
E: 3
44
N: 3
E: 3
45
N: 3
E: 3
46
N: 3
E: 3
47
N: 3
E: 3
48
N: 3
E: 3
49
N: 3
E: 3
50
N: 3
E: 3
51
N: 3
E: 3
52
N: 3
E: 3
53
N: 3
E: 3
54
N: 3
E: 3
55
N: 3
E: 3
56
N: 3
E: 3
57
N: 3
E: 3
58
N: 3
E: 3
59
N: 3
E: 3

**c.**

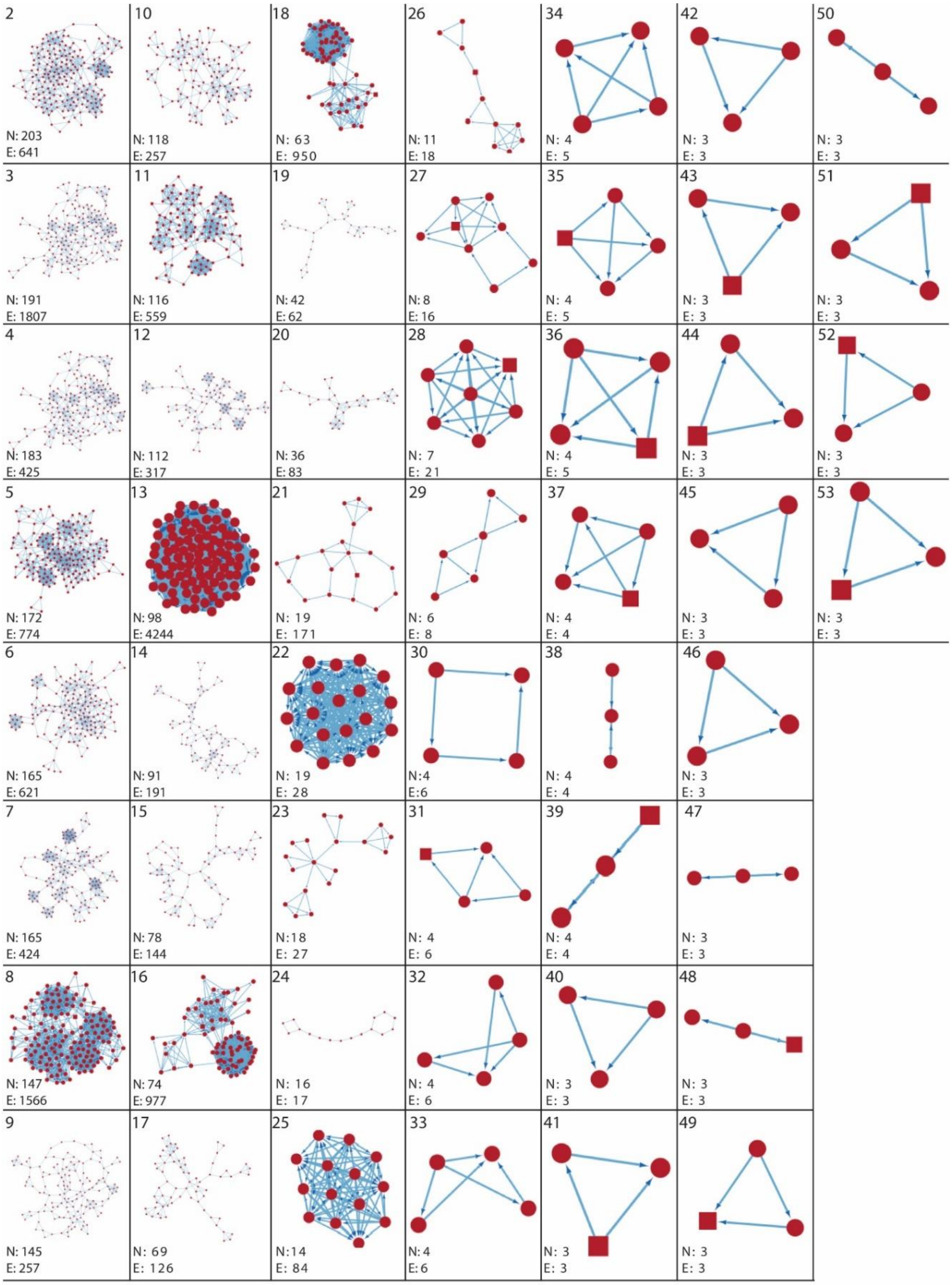

**Fig. S2.** We constructed the network and retrieved node degree, betweenness, and closeness centrality scores using Cytoscape (v3.10.0) StringApp (v2.0.1) [1-3]. Molecular Complex Detection (MCODE)_v2.0.3 method [4] was used to identify the molecular clusters of the entire network. MCODE used the graph theoretic clustering algorithm to identify densely connected regions in a large protein-protein interaction network (PPIn) representing molecular complexes [4]. We also investigated the functional roles of oxidative, cellular, and antibiotic stress response genes through ClueGO (v2.5.10), a Cytoscape plug-in [2, 5]. We used default parameter values during network construction via the String App (i.e., Network type: full String network, Confidence cutoff: 0.4) and selective parameter values during MCODE analysis like, Find Clusters: in Whole Network; Network Scoring - a) Include Loops: Turn off, b) Degree Cutoff: 10; Cluster Finding- a) Haircut: Turn on, b) Fluff: Turn off, c) Node Score Cutoff: 0.2, d) K-Core: 2, e) Max. Depth: 100. During ClueGO analysis, default parameter values were used, such as Analysis Mode: Functional Analysis, Visual Style: Groups, Ontologies: Biological and cellular process. **a.** MCODE generated size-based clusters at 3 h Amp treatment. N and E refer to nodes and edges, respectively. A total of 62 clusters were detected, and here, we present 61 clusters arranged in size-based ranking order. **b.** MCODE generated size-based clusters at 6 h Amp treatment. N and E refer to nodes and edges, respectively. A total of 59 clusters were detected, and here, we present 58 clusters arranged in size-based ranking order. **c.** MCODE generated size-based clusters at 24 h Amp treatment. N and E refer to nodes and edges, respectively. A total of 54 clusters were detected, and here, we present 53 clusters arranged in size-based ranking order.

**a.**

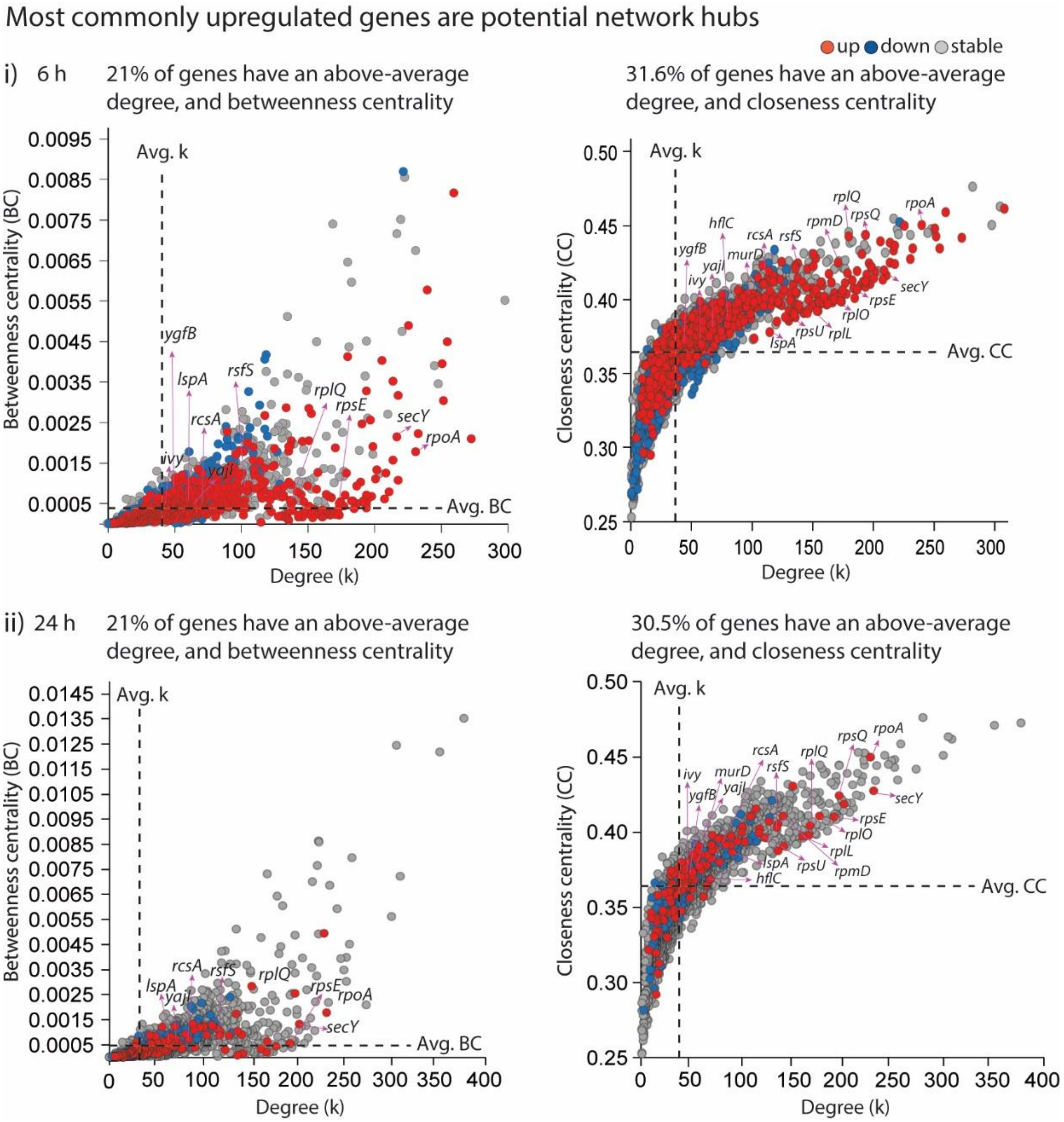
Most commonly upregulated genes are potential network hubs
up
down
stable
i) 6 h
21% of genes have an above-average degree, and betweenness centrality
31.6% of genes have an above-average degree, and closeness centrality
Avg. k
Betweenness centrality (BC)
Avg. BC
Degree (k)
Closeness centrality (CC)
Avg. CC
ygfB
lspA
rsfS
rcsA
rplQ
rpsE
secY
rpoA
ivy
yajI
hflC
murD
rpmD
rpsQ
rpsU
rplL
rplO
ii) 24 h
21% of genes have an above-average degree, and betweenness centrality
30.5% of genes have an above-average degree, and closeness centrality

**b.**

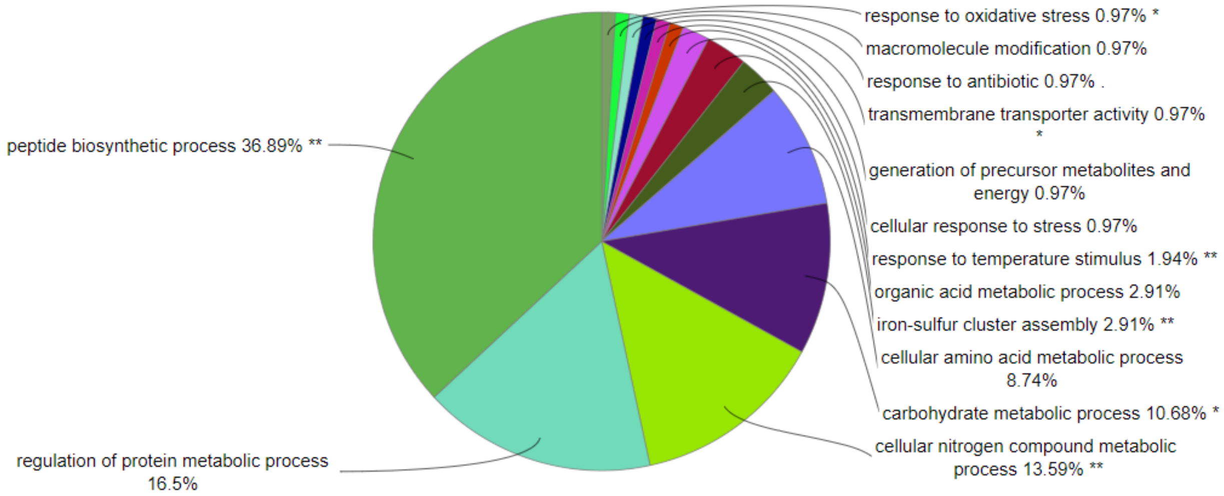
response to oxidative stress 0.97% *
macromolecule modification 0.97%
response to antibiotic 0.97% .
transmembrane transporter activity 0.97% *
generation of precursor metabolites and energy 0.97%
cellular response to stress 0.97%
response to temperature stimulus 1.94% **
organic acid metabolic process 2.91%
iron-sulfur cluster assembly 2.91% **
cellular amino acid metabolic process 8.74%
carbohydrate metabolic process 10.68% *
cellular nitrogen compound metabolic process 13.59% **
peptide biosynthetic process 36.89% **
regulation of protein metabolic process 16.5%

**c.**

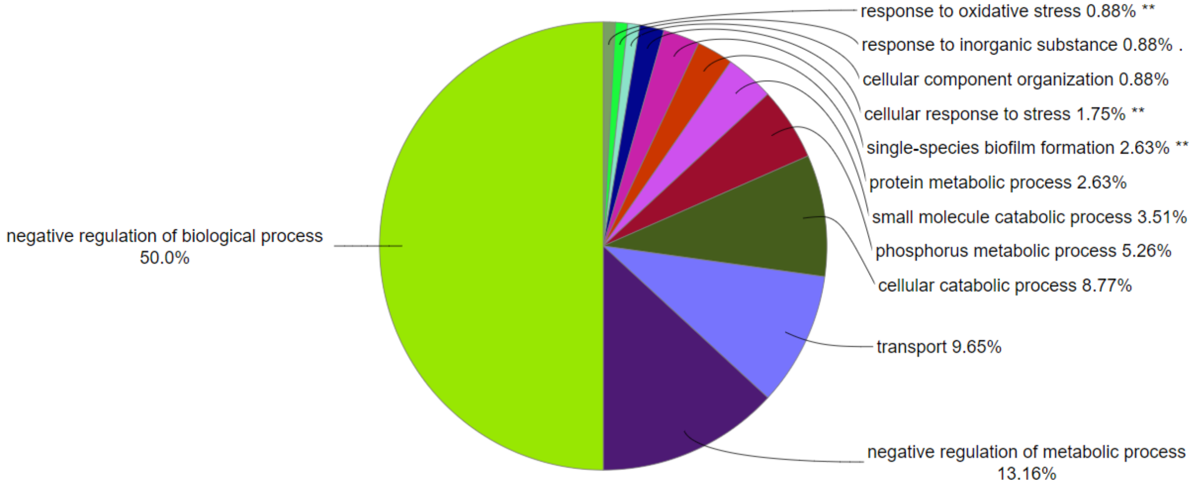
response to oxidative stress 0.88% **
response to inorganic substance 0.88% .
cellular component organization 0.88%
cellular response to stress 1.75% **
single-species biofilm formation 2.63% **
protein metabolic process 2.63%
small molecule catabolic process 3.51%
phosphorus metabolic process 5.26%
cellular catabolic process 8.77%
transport 9.65%
negative regulation of metabolic process 13.16%
negative regulation of biological process 50.0%

**d.**

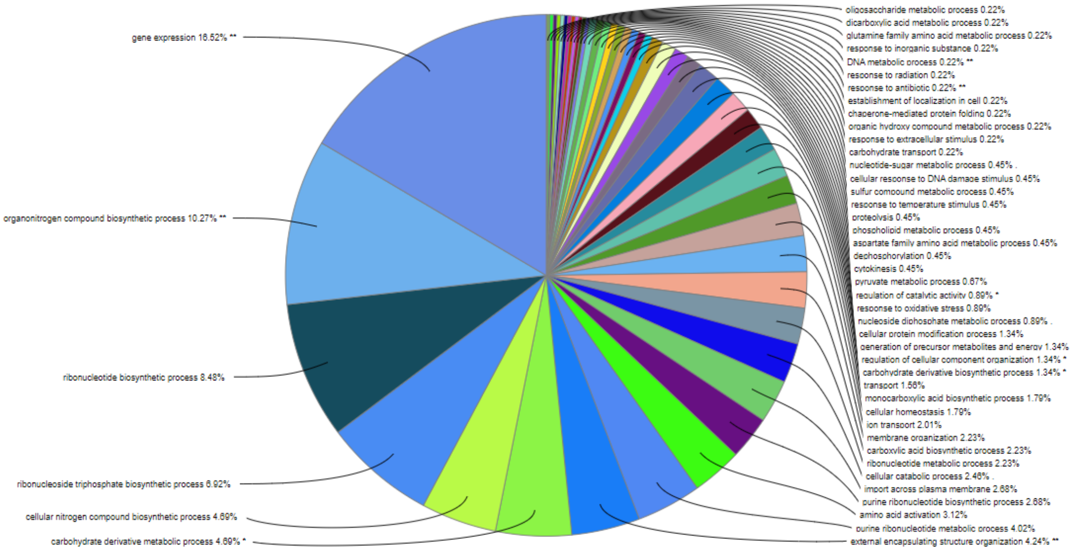
gene expression 16.52% **
organonitrogen compound biosynthetic process 10.27% **
ribonucleotide biosynthetic process 8.48%
ribonucleoside triphosphate biosynthetic process 6.92%
cellular nitrogen compound biosynthetic process 4.69%
carbohydrate derivative metabolic process 4.69% *
oligosaccharide metabolic process 0.22%
dicarboxylic acid metabolic process 0.22%
glutamine family amino acid metabolic process 0.22%
response to inorganic substance 0.22%
DNA metabolic process 0.22% **
response to radiation 0.22%
response to antibiotic 0.22% **
establishment of localization in cell 0.22%
chaperone-mediated protein folding 0.22%
organic hydroxy compound metabolic process 0.22%
response to extracellular stimulus 0.22%
carbohydrate transport 0.22%
nucleotide-sugar metabolic process 0.45% .
cellular response to DNA damage stimulus 0.45%
sulfur compound metabolic process 0.45%
response to temperature stimulus 0.45%
proteolysis 0.45%
phospholipid metabolic process 0.45%
aspartate family amino acid metabolic process 0.45%
dephosphorylation 0.45%
cytokinesis 0.45%
pyruvate metabolic process 0.67%
regulation of catalytic activity 0.89% *
response to oxidative stress 0.89%
nucleoside diphosphate metabolic process 0.89% .
cellular protein modification process 1.34%
generation of precursor metabolites and energy 1.34%
regulation of cellular component organization 1.34% *
carbohydrate derivative biosynthetic process 1.34% *
transport 1.56%
monocarboxylic acid biosynthetic process 1.79%
cellular homeostasis 1.79%
ion transport 2.01%
membrane organization 2.23%
carboxylic acid biosynthetic process 2.23%
ribonucleotide metabolic process 2.23%
cellular catabolic process 2.46% .
import across plasma membrane 2.68%
purine ribonucleotide biosynthetic process 2.68%
amino acid activation 3.12%
purine ribonucleotide metabolic process 4.02%
external encapsulating structure organization 4.24% **

**e.**

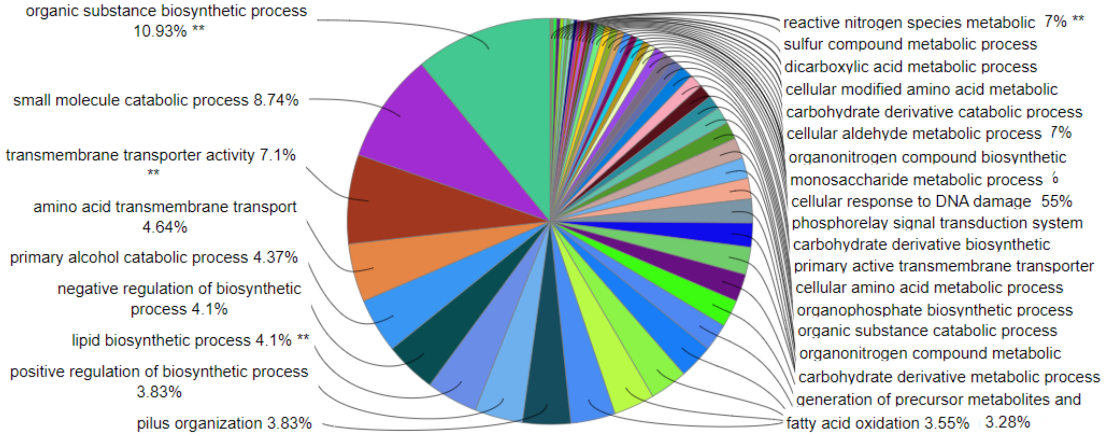
organic substance biosynthetic process 10.93% **
small molecule catabolic process 8.74%
transmembrane transporter activity 7.1% **
amino acid transmembrane transport 4.64%
primary alcohol catabolic process 4.37%
negative regulation of biosynthetic process 4.1%
lipid biosynthetic process 4.1% **
positive regulation of biosynthetic process 3.83%
pilus organization 3.83%
reactive nitrogen species metabolic 7% **
sulfur compound metabolic process
dicarboxylic acid metabolic process
cellular modified amino acid metabolic
carbohydrate derivative catabolic process
cellular aldehyde metabolic process 7%
organonitrogen compound biosynthetic
monosaccharide metabolic process
cellular response to DNA damage 55%
phosphorelay signal transduction system
carbohydrate derivative biosynthetic
primary active transmembrane transporter
cellular amino acid metabolic process
organophosphate biosynthetic process
organic substance catabolic process
organonitrogen compound metabolic
carbohydrate derivative metabolic process
generation of precursor metabolites and
fatty acid oxidation 3.55% 3.28%

**f.**

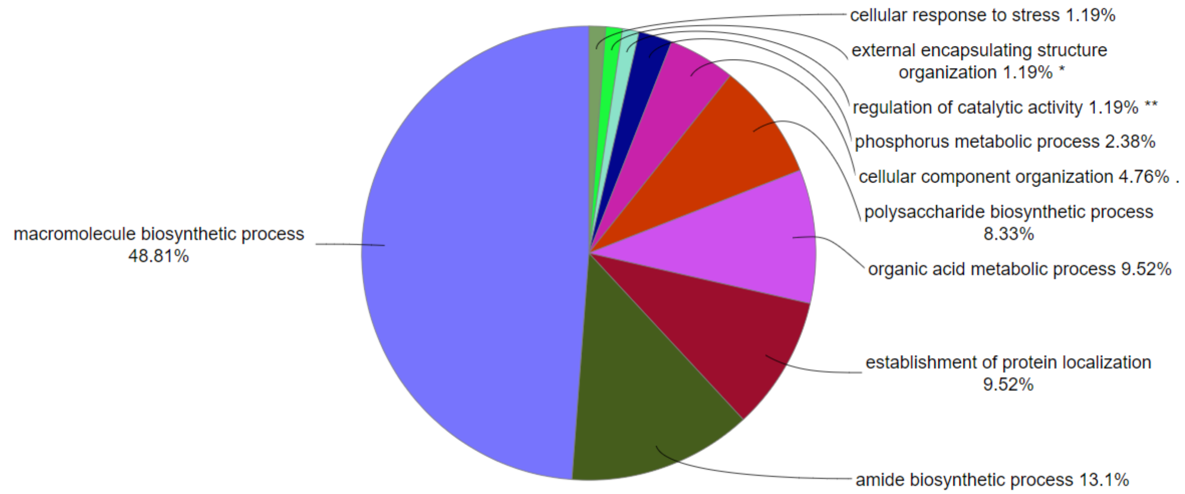

**g.**

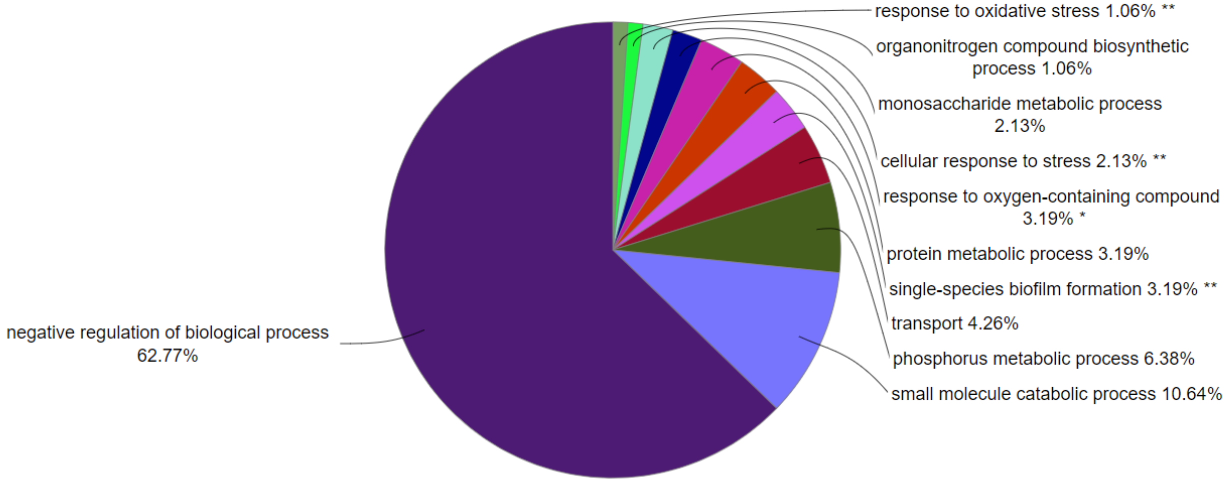

**Fig. S3. a.** Most upregulated genes are potential network hubs. Identification of potential network hubs: Degree (k) vs. Betweenness Centrality (BC) and Degree (k) vs. Closeness Centrality (BC) analyses identified commonly upregulated genes as potential network hubs during Amp treatment over time. **i.** At 6 h, a substantial number of genes (21%) showed above-average centrality scores in the k vs. BC analysis, while 31.6% of genes displayed above-average centrality scores in the k vs. CC analysis. **ii.** Similarly, at 24 h, 21% of genes showed above-average centrality scores in the k vs. BC analysis, with 30.5% of genes showing above-average centrality scores in the k vs. CC analysis. These findings highlight the pivotal roles played by these genes within the network and underscore their potential as key targets for further investigation. **b.** 3 h upregulated gene functions and statistics. **c**. 3 h downregulated gene functions and statistics. **d.** 6 h upregulated gene functions and statistics. **e.** 6 h downregulated gene functions and statistics. **f.** 24 h upregulated gene functions and statistics. **g.** 24 h downregulated gene functions and statistics. **Group p-value Corrected with Bonferroni step-down.

**Table S1. a.** Commonly upregulated genes at 3 h, 6 h, and 24 h Amp treatment compared to 0 h untreated. **b.** Upregulated or downregulated genes at 3, h, 6 h, and 24 h compared to compared to 0 h untreated.

| **a.** | |
|---|---|
| **Gene** | **Functions** |
| *acrB* | Multidrug efflux pump RND permease AcrB |
| *bioF* | Play a role in biofilm formation |
| *hflC* | Regulator of FtsH protease |
| *iscX* | Accessory iron-sulfur cluster assembly protein IscX |
| *ivy* | Periplasmic chaperone, inhibitor of vertebrate C-type lysozyme |
| *lspA* | Lipoprotein signal peptidase |
| *murD* | Cell wall formation (probable) |
| *rplL* | 50S ribosomal subunit protein L7/L12 dimer |
| *rplO* | 50S ribosomal subunit protein L15 |
| *rplQ* | 50S ribosomal subunit protein L17 |
| *rpmD* | 50S ribosomal subunit protein L30 |
| *rpoA* | RNA polymerase subunit α |
| *rpsE* | 30S ribosomal subunit protein S5 |
| *rpsQ* | rRNA binding proteins; plays a role in translational accuracy |
| *rpsU* | 30S ribosomal subunit protein S21 |
| *secY* | Required for efficient protein translocation across the cytoplasmic membrane |
| *ycfJ* | Uncharacterized protein; plays role in biofilm formation |
| *ygfB* | Uncharacterized lipoprotein |
| *yajI* | Uncharacterized lipoprotein |
| *yciF* | Uncharacterized protein; upregulated when bacteria experience stress conditions and is highly conserved in a range of bacterial species |
| *yciY* | Uncharacterized protein |
| *yfgD* | Uncharacterized protein |
| *ykiA* | Putative uncharacterized protein |
| *ypeB* | Uncharacterized protein |
| *rcsA* | Component of the Rcs signaling system, which controls transcription of numerous genes |
| *wcaB* | Involved in the pathway of slime polysaccharide biosynthesis, which is part of Slime biogenesis. |
| *rsfS* | Ribosomal silencing factor RsfS |

| **b.** | | | | | |
|---|---|---|---|---|---|
| | **Expression** | | **Amp** | | **Cip** |
| **Gene** | **Function** | **3 h** | **6 h** | **24 h** | **6 h** |
| *oxyR* | Antioxidative defense pathway | Stable | Up | Stable | up |
| *dnaK* | Global regulator | Up | Stable | Stable | stable |
| *clpB* | Global regulator | Up | Stable | Stable | stable |
| *rpoS* | Global regulator | Stable | Up | Stable | stable |
| *sucB* | Energy production | Up | Stable | Stable | stable |
| *relA* | Stringent response | Stable | Stable | Stable | stable |
| *mqsR* | Toxin-antitoxin system | Down | Down | Down | down |
| *recA* | SOS response | Stable | Up | Stable | up |
| *lon* | TA module/protease | Up | Stable | Stable | stable |
| *relE* | TA module | Down | Down | Down | down |
| *hipA* | TA module/HipA toxin functions as a serine/threonine kinase that inhibits cell growth, | Stable | Stable | Stable | stable |
| *dinJ* | TA module | Down | Down | Down | down |
| *phoU* | Global Regulator | Stable | Up | Stable | stable |
| *smpB* | Trans-translation | Stable | Stable | Stable | up |
| *glpD* | Energy production | Stable | Down | Down | stable |
| *umuD* | SOS response | Stable | Down | Stable | up |
| *tnaA* | Signaling pathway | Stable | Stable | Stable | down |

| | | | | | |
|---|---|---|---|---|---|
| *pspF* | Signaling pathway | Stable | Stable | Stable | stable |
| *uvrA* | SOS response | Stable | Stable | Stable | stable |

**Table S2.** A large fraction of genes are neither downregulated nor upregulated when comparing the persister state to the stationary state during Amp treatment over time (using an adjusted p-value of <0.1 and 2-fold change as a cutoff). *E. coli* Dh5alphaZ1 has 4357 annotated genes.

| **Antibiotic treatment time** | **3 h** | **6 h** | **24 h** |
|---|---|---|---|
| Genes with unchanged expression | 3775 | 2773 | 3941 |
| % genes with unchanged expression | 86% | 63% | 90% |

**Table S3.** The number of upregulated and downregulated genes related to critical functions changes during Amp treatment over time.

| | **No. of genes at 3 h** | | **No. of genes at 6 h** | | **No. of genes at 24 h** | |
|---|---|---|---|---|---|---|
| **Functions** | **Upregulated** | **Downregulated** | **Upregulated** | **Downregulated** | **Upregulated** | **Downregulated** |
| Response to oxidative stress | 11 | 13 | 14 | 19 | 0 | 12 |
| Response to antibiotic | 10 | 0 | 34 | 18 | 0 | 0 |
| Cellular response to stress | 17 | 31 | 48 | 82 | 11 | 24 |

**Table S5. Strains and plasmids.** The cultures were grown in MMB+, as described in our previous work [6, 7]. We used 0.5% glycerol as the carbon source and each of the 20 canonical amino acids (0.04 mg/mL) for fast-growing cultures. Cultures were grown in MMB+ or on Miller's lysogeny broth (LB) agar plates with antibiotics to maintain the plasmid. All cultures were incubated at 37 °C and shaken at 250 or 300 rpm.

| **Strain and function** | **Plasmid and function** |
|---|---|
| *Escherichia coli* DH5αZ1 is a derivative of *E. coli* K12 strain, and it has been used in our previous persistence studies [6, 8]. | Plasmid p24KmNB82 plasmid was used for all persister assays, and kanamycin (Km, 0.025 mg/mL) was used to maintain the plasmid in *E. coli*. |

**Table S6. RNA sample preparation, sequencing, and analysis.** RNA harvest and analysis for **a. s**tationary phase cells as untreated (0 h stationary phase) and **b.** log phase as untreated (0 h log) were different

| **a.** | |
|---|---|
| **Isolation Steps** | **Harvested RNA from persisters using our previously established methodology and incorporated previously published data (NCBI GEO: GSE156896) Ref [9]** |
| 1 | Cultures were first grown to stationary phase before starting antibiotic treatment. |
| 2 | Cells were treated with lethal concentrations of Amp (0.1 mg/ml) or Cip (0.001 mg/ml). |
| 3 | Samples were treated with 0.2% Tween and 0.005 mg/ml chloramphenicol, then immediately filtered out of the solution using rapid filtration. |
| 4 | Cells were resuspended in lysis buffer containing 25 mM Tris pH 8.0, 25 mM $NH_4Cl$, 10 mM MgOAc, 0.8% Triton X-100, 0.1 U/µL RNase-free DNase I, 1 U/µL Superase-In, ~ 60 U/µl ReadyLyse Lysozyme, 1.55 mM Chloramphenicol, and 17 µM 5′-guanylyl imidodiphosphate (GMPPNP). |
| 5 | The solution was then frozen and stored at -80 °C. |
| 6 | Frozen pellets were thawed on ice and centrifuged for 20 min at 16,000 ×*g* at 4 °C with 0.3% Sodium Deoxycholate to pellet particulate matter. |
| 7 | The supernatant was removed, and |
| 8 | RNA was column purified (Qiagen miRNeasy; 217004), |
| 9 | followed by rRNA removal and cDNA library preparation (NEBNext Ultra II; E7103S). |
| 10 | Libraries were multiplexed, pooled, and sequenced using Illumina HiSeq paired-end sequencing by GENEWIZ. |
| **Analysis** | Transcriptomes were compared to an untreated population (stationary phase, 0 h) similarly to Ref. [9] |

| **b.** | **Procedure** |
|---|---|
| **Isolation Steps** | Similar to **a.** with two changes: cells were grown to mid-log phase (~OD 0.5) rather than stationary phase before antibiotic treatment, and (2) RNA was harvested by pelleting the cells and removing the supernatant. |
| **Analysis steps** | |
| 1 | FastQC was used to assess the quality of RNA-seq data. |
| 2. | Raw counts for the samples were generated using a combination of Geneious Prime NGS software [10] and custom Python scripts adapted from our previous transcriptome analyses [11]. |
| 3. | Differential gene expression analysis was performed using the DESeq2[12], edgeR [13], and Limma [14] R packages. TMM normalization was applied using edgeR calcNormFactors function with the default LogratioTrim set to 0.3. Cyclic Loess normalization was applied using the normalizeCyclicLoess function from the Limma package. |
| 4. | Significant changes in gene expression were defined as greater than 4-fold with an adjusted p-value <0.1 for stationary-growth phase data and <0.05 for log growth phase data. |
| 5. | Venn diagrams was generated using InteractiVenn [15]. |
